# Evolution is not uniform along protein sequences

**DOI:** 10.1101/2022.04.30.490124

**Authors:** Raphaël Bricout, Dominique Weil, David Stroebel, Auguste Genovesio, Hugues Roest Crollius

## Abstract

Amino acids evolve at different speeds within protein sequences, because their functional and structural roles are different. However, the position of an amino-acid within the sequence is not known to influence this evolutionary speed. Here we discovered that amino-acid evolve almost twice faster at protein termini than in their centre, hinting at a strong topological bias along the sequence length. We further show that the distribution of functional domains and of solvent-accessible residues in proteins readily explain how functional constrains are weaker at their termini, leading to the observed excess of amino-acid substitutions. Finally, we show that methods inferring sites under positive selection are strongly biased towards protein termini, suggesting that they may confound positive selection with weak negative selection. These results suggest that accounting for positional information should improve evolutionary models.

## Main text

Rates of evolution vary greatly between protein-coding gene families, for example in correlation with their expression levels (*1, 2*), their function (*3*) or with translational selection (*4*) effects. Within a given gene family, molecular rates of evolution can also vary within lineages (*5*) and between lineages (*6*). Within proteins themselves, rate heterogeneity among amino-acid sites is influenced by their implication in functional domains and by structural constraints in the folded protein (*7*). Accounting for such heterogeneity in evolutionary models is critical to accurately infer phylogenies and estimate cases of positive selection, and elaborate models have been developed to achieve this (*8*), generally by estimating site-specific rates in a maximum likelihood framework employing Markov models of sequence evolution (*9*–*11*).

Surprisingly, the impact of the position of a given amino-acid in the sequence relative to the protein start and end on the rate of molecular evolution has not yet been investigated. To address this question, we measured the rates of amino-acid changing substitutions (non-synonymous, dN) and silent substitutions (synonymous, dS) at individual codons positions and average them over thousands of CDS sequences. Specifically, we computed multiple sequence alignments (MSA) of CDS from 16,810 primate gene families to identify fixed mutations (substitutions, insertions and deletions) that took place in these sequences during the evolution of 26 primate species. The results show a strong excess of such changes towards the sequence extremities (Fig. 1A), leading to a distinctive U-shaped pattern. Looking further into substitutions at each individual codon position, we computed position-specific codon average evolutionary rates (Fig. S1) to examine separately the dN and the dS. While the dS remains remarkably constant along the CDS length (average dS=0.052), the dN increases significantly in the region spanning the first and last 50 codons (Fig. 1B). We observe a similar bias when computing dN and dS along the CDS of 7,513 plant (Fabids) gene families, which were subjected to an approximately 8-fold higher divergence rate than primates (Fig. 1C). In summary, the dN appears to be driving the distinctive U-shaped pattern of total substitutions and dN/dS in gene coding sequences (Fig. 1D).

**Fig. 1.**
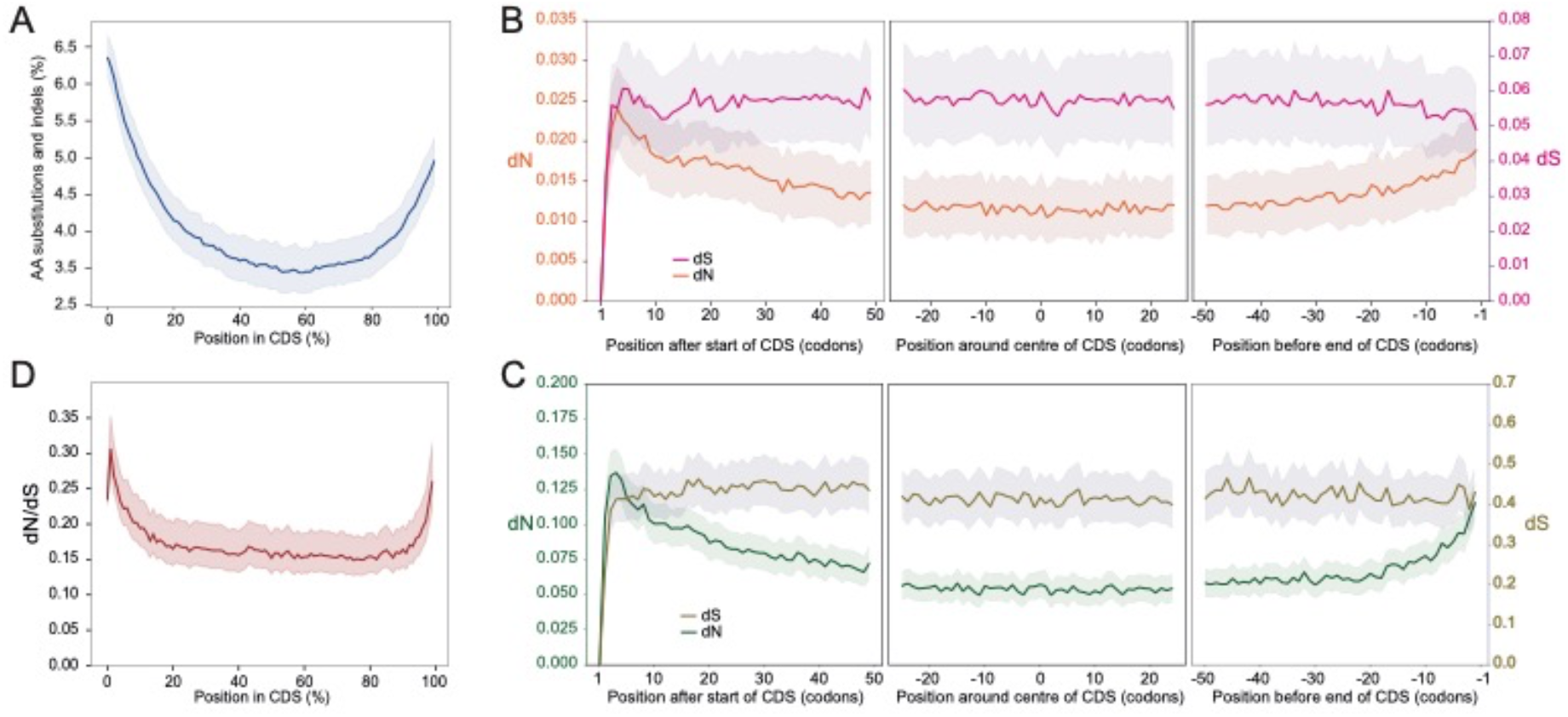
(**A**). Frequency of amino acid substitutions, insertion and deletions computed in 16,810 primate multiple sequence (CDS) alignments (MSA), rescaled from 0 to 100% of the coding sequence length. (**B**). Distribution of silent (dS) and non-synonymous (dN) substitution rates computed at each codon position from random pairs of sequences sampled from 16,248 primate MSA without alignment gaps and shown here for the first 50 codons (left panel), the central 50 codons (middle panel) and the last 50 codons (right panel). (**C**). Same as in B but for pairs of CDS sampled from 7,513 plant MSA. (**D**). The distribution of dN/dS ratio for dN and dS values shown in B but across the entire CDS length rescaled from 0 to 100%. In all panels the shaded the area represents the 95% confidence interval.

Computing substitutions in a multiple alignment of coding sequences is a multi-step process, with many potential sources of technical biases which could potentially explain this pattern (*12, 13*). We conducted a series of experiments to exclude annotation errors, multiple alignment artefacts and compositional biases (Supplementary material, Fig. S2), showing that our observations are robust to controls designed to address possible technical artefacts in the process from CDS annotation to substitution calculations.

We next examined biological or evolutionary explanations. We eliminated the possibility that a stronger mutation rate at CDS extremities would fuel the increased dN because the dS, which would be much more sensitive to the mutation rate, is essentially constant along CDS length (Fig. 1B,C). We next reasoned that a weaker negative selection at protein termini might be caused by weaker functional constraints. Predicted protein domains capture a large fraction of amino acids involved in structural and functional roles in protein sequences, and their prediction relies on sequence similarity and structural information (*14*). Both of these features make them good proxies for sites under evolutionary constraints. We computed the distribution of protein domains predicted by different methods along the 12,067 human protein sequences involved in our set of primate gene families, and we show that domains are strongly depleted at protein edges (Fig. 2A and fig. S3A-C). This dome-like shape is caused by the lower probability of domains overlapping amino acids immediately adjacent to these termini, since they cannot physically overlap the termini themselves. The distribution of domains decreases sharply towards protein edges regardless of the length of the protein sequences (Fig. S3D-E), supporting a scenario where all proteins are similarly affected by a deficit of domain-induced negative constraints at their edges. The dome-shaped distribution of domains mirrors the distinctive U-shaped distribution of the dN/dS ratio (Fig. 1A), consistent with our initial hypothesis that a depletion of domains at the edges of proteins would make them more permissive to non-synonymous changes and indels because of weaker selective constraints. To test this more directly, we distinguished codons that code for amino acids involved in a domain from those that do not, and computed the dN/dS for each category separately (Fig. 2B). In line with the above expectation, the dN/dS bias disappears when computed exclusively inside domains. Again, the difference in dN/dS behaviour is largely caused by the dN, since the dS remains constant both inside and outside of domains and is almost identical in both categories throughout the protein lengths (Fig. S4). These results strongly support a model in which selective constraints are significantly weaker at protein edges.

**Fig. 2.**
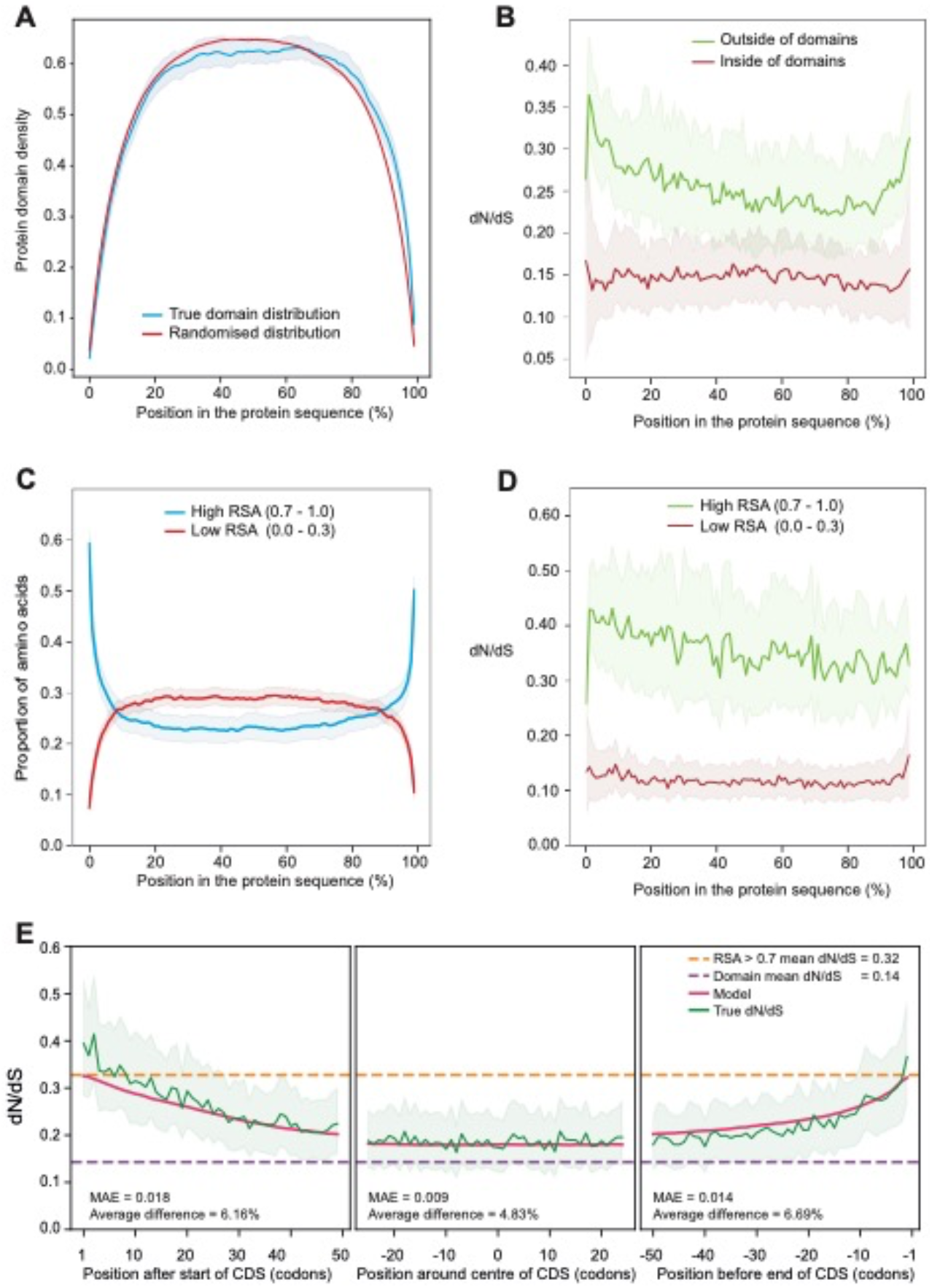
(**A**) The distribution of protein domains from the PFAM database in 12,067 human proteins rescaled from 0-100% of their length (blue line). The red line shows the distribution of the same domains in random non-overlapping positions in the same sequences. (**B**) dN/dS computed in 16,248 alignments of at least 2 CDS sequences from 26 primate genomes, where sites inside (dark red line) and outside (green line) PFAM domains are distinguished. (**C**) The distribution of frequency of amino acids with high (blue line) and low (red line) Relative Solvent Accessibility (RSA) computed by pCASA on 3D structure predicted by AlphaFold on 23,391 human protein sequences, rescaled to 0-100% of the length. (**D**) dN/dS computed on 7,614 sequences common to the AlphaFold and Ensembl primate CDS datasets, where sites with high (green line) and low (dark red line) RSA are distinguished. (**E**) Proposed model were the mean dN/dS in domains and in highly accessible regions (RSA > 0.7) are weighted according respectively to the percent of codons overlapping domains and residues with RSA > 0.7. The Mean Absolute Error (MAE) and percent average error are indicated for each panel. In all panels the shaded the area represents the 95% confidence interval.

We next investigated how structural constrains, or lack thereof, may also influence evolutionary rates at protein termini. A protein sequence is folded in space through both local and distant amino-acid interactions. Amino acids which are free of those interactions are conversely accessible to solvents, and typically found on the surface of the folded protein. Because such structural interactions are linked to the function of a protein, it is well-established that evolutionary rates differ between residues depending on their solvent accessibility(*15*–*17*). The precise relationship between solvent accessibility, evolutionary rates and amino-acid position along the sequence has however never been ascertained, mostly because N- and C-terminal regions of proteins are often removed to facilitate crystallisation prior to structure solving, or are generally poorly resolved in electron density maps. To circumvent this issue, we analysed 23,391 structures predicted by Alphafold (*18*) on complete human protein sequence to compute the relative solvent accessibility (RSA) of each residue (Fig. S5A). We note that RSA values follow a bimodal distribution with a distinctive peak above 0.7 depleted in residues included in protein domains. Conversely, residues with RSA below 0.3 are enriched in functional domains. We took residues from these two extremes categories to compute their distribution along protein sequences, and we find that solvent accessibility increases sharply at protein termini, consistent with weak structural constraints in these regions (Fig. 2C). Critically, the dN/dS rate is low and constant along protein length in sites with low accessibility, while it is high in highly accessible regions (Fig. 2D). In both categories, the marked increase in dN/dS at sequence extremities shown in Figure 1 is absent, indicating that solvent accessibility is likely a strong marker of the decrease in selective pressure observed in the N- and C-terminal region of proteins. Because both functional domains and RSA seem to drive dN/dS variation at protein termini, we applied a model where the average dN/dS at a given residue is the sum of the average dN/dS in domains and in high RSA (RSA > 0.7), each weighted by the proportion of residues in their category (Supplementary Material). The model reproduces the observed dN/dS with remarkable accuracy in human proteins (mean difference = 5.9%, Fig. 2E), suggesting that functional domains and RSA are two variable that are sufficient to explain the bias in average dN/dS along proteins sequences. The slight asymmetry in dN/dS bias between N- and C-termini observed in the model, reminiscent of the asymmetry observed in Figure 1B, largely disappears when we remove the 15.8% of proteins labelled with a signal peptide at the N-terminus (Fig. S5B). Signal peptide, which are known to be highly variable in sequence (*19*) are therefore likely to provide additional relief from the evolutionary pressure measured in this region.

The dN/dS bias at protein ends is measurable by averaging thousands of sites at any given position. Is it also significant at the level of individual sequence alignments? This is important if evolutionary models applied to single gene families are likely to be affected. To address this, we computed a correlation between codon position and dN/dS for 12,322 multiple sequence alignments, separately for the 50 codons at beginning and at the end of coding sequences (Fig. 3A).

**Figure 3.**
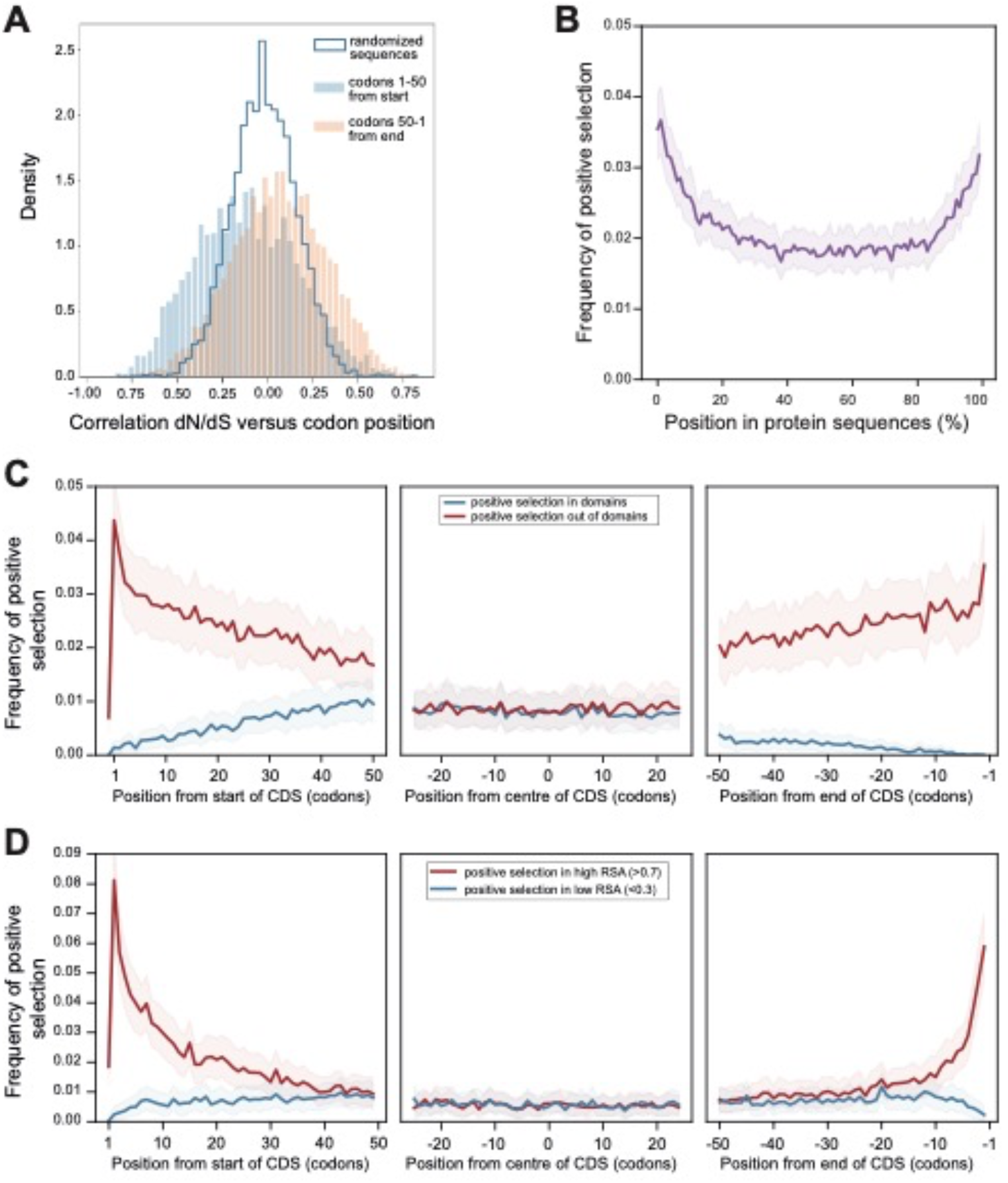
(**A**) Distribution of Pearson correlation coefficients between dN/dS values and their position in mouse CDS from 12,322 alignments of 5 to 20 rodent sequences. Positions are either the first 50 (filled blue bars) or the last 50 (filled pink bars) codons. The blue line shows the distribution for the first and last 50 codons in the same mouse sequences but where their positions were randomized. (**B**) Distribution of the frequency of sites under positive selection in 12,170 rodent CDS rescaled to 0-100% of their length (**C**) Distribution of the frequency of sites under positive selection in the first, middle and last 50 codons of 12,170 rodent CDS, distinguishing sites inside (blue line) and outside (red line) PFAM domains. (**D**) Distribution of the frequency of sites under positive selection in the first, middle and last 50 codons of 5,893 rodent CDS, distinguishing sites with high (red line) and low (blue line) RSA.

Compared to a control where amino-acid positions are randomised, the distribution of correlations are significantly shifted towards negative (p-value=8.10^−80^; t-test) and positive (p-value = 6.10^−46^; t-test) values for the start and end regions of coding sequences, respectively. This reflects the existence of a measurable increase in dN/dS towards CDS edges, even in individual sequences. We propose that this pattern is caused by the same factors as for the average sequences analysed previously (Fig. 1B), respectively the biased domain distribution and solvent accessibility.

These results immediately raise questions for the identification of positive selection in proteins sequences, because a significantly elevated dN compared to some background rate is generally taken as evidence of adaptive changes (*20*). The U-shaped bias in dN observed in our study suggests that relaxation of constraints at protein edges might confound tests of positive selection. To investigate this, we estimated sites under positive selection in a set of 12,170 Rodent gene trees using a site model (methods). We found that sites estimated under positive selection are strikingly enriched at protein edges (Fig. 3B), and that this enrichment is specifically attributed to sites located outside of functional domains (Fig. 3C) and to residues with high solvent accessibility (Figure 3D). Notably, the same bias towards coding sequence extremities can be observed in several recent published scans for positive selection (Figure S6). Interestingly, while this bias is conspicuous for sites estimated to have been subject to positive selection using bioinformatic methods, it is not the case for experimentally verified sites, although our compilation of cases for this category is too small to draw general conclusions.

We reveal a pattern of evolutionary rate along coding sequences that has so far remained concealed: the average amino acid substitution rate (dN) increases towards the extremities of the sequence. We found this pattern by assembling observations that were sometimes known quantitively or intuitively in the field but never connected with respect to codon or amino-acid positions in sequences. This patterns provide insights into the elusive mechanism driving evolutionary rate heterogeneities (*7*). First noted by Perutz and colleagues on haemoglobin (*21*) and confirmed by many studies since, protein surfaces evolve faster than their interior, where structural constraints, residue interactions and functional sites are most enriched and solvent accessibility is very low (*15, 16*). Attempts at explaining evolutionary rate heterogeneity have thus mainly focused on this paradigm, that structural constraints governed by complex spatial interactions create a range of selective pressures on amino acids, but these are still hard to predict from the sequence itself.

We note that in the present study molecular rates are not dependent on a substitution model, as they do not rely on ancestral state inferences in coding sequences, and molecular rates are computed on MSA without gaps, which are known to introduce biases. Also, our finding that the average dS is constant along CDS length (Figs. 1B, 1C) should not be interpreted as meaning that dS does not vary or is not subject to site heterogeneity in individual genes, as this has been shown in many studies (*22*).

If protein termini are under lower evolutionary pressure, why are they not cropped by micro-deletions in the course of evolution? The example of signal peptides, but also of amino-acids carrying specific epigenetic marks (e.g. methyl or acetyl groups) in histones, illustrate why maintaining structurally flexible regions may be functionally important.

Methods designed to identify positive selection are sensitive to false positives, potentially caused by factors such as variable effective population size (*23*), biased gene-conversion (*24*), multi-nucleotide mutations (*25*) and punctual relaxation of selective pressure in a lineage (*5, 26*). Here we show that sites inferred as having experienced a period of positive selection are conspicuously enriched in regions with high dN caused by low selective pressure, suggesting that they may contain a high proportion of false positives. This is consistent with the observation that experimentally tested positively selected sites are, on the contrary, depleted at sequence extremities. Of note, we observed that the synonymous rate dS is constant along protein length, thus providing little leverage for background model adjustments to counteract this effect in statistical tests of positive selection. Considering the excess of positive selection inferences at protein extremities as false positives would also be consistent with expectations that selection for advantageous traits would operate predominantly where functional domains and structural constraints are most frequent, i.e. away from the extremities (*27*). It has also been previously shown that non-adaptive changes as well as positively selected sites are significantly enriched on the surface of proteins where solvent accessibility is high, emphasizing the difficulty in distinguishing them in these regions (*17*). Altogether, we propose that intervals between functional domains display a neutrally evolving size and weaker structural constraints, largely causing lower selective pressure at protein termini. Accounting for this bias in models of molecular evolution should improve their handling of site heterogeneity and accuracy of adaptive evolution inference.

## Acknowledgements

We thank P. Vincens and the informatics service at IBENS for support, and Alexandra Louis, Guillaume Louvel, François Giudiucelli and Nicolas Lartillot for helpful discussions.

## Author contributions

R.B., A.G. and H.R.C. conceived the study and analysed results. D. W. and D.S. analysed results. All authors contributed to the writing of the manuscript.

## Data and material availability

The scripts and data necessary to generate all figures, including in supplementary Information, have been deposited in Zenodo (doi:10.5281/zenodo.6472876).

## Supplementary Materials

*Material and Methods Fig S1 to S6*

*Table S1*

*References*

