## Supplementray Material for "Evolution is not uniform along protein sequences"

### Evolution is not uniform along protein sequence length

***This PDF file includes:***

*Supplementary Results*
*Material and Methods*
*Fig. S1 to S6*
*Table S1*
*References*

**Supplementary Results**

##### 1. Controlling for experimental biases

First, the annotation of protein coding genes in genomic sequences is subject to many incorrect gene model predictions, especially in non-model organisms. Although all CDS used in this study possess an ATG start codon and a stop codon, more frequent errors at the 5' and 3' ends of a CDS may lead to imprecise definitions of the start and stop codons and inclusion of non-coding base triplets as codons, which would lead to an increased dN values in these regions. To circumvent such potential annotation errors, we measured amino-acid substitutions between human and mouse coding sequences annotated directly in sequenced transcripts, not in genomic DNA, and show that the substitution excess at the 5' and 3' extremities of the CDS is similar in intensity to the pattern observed in genome-based CDS annotations (Fig. S2A). In contrast, if we force a dN/dS computation in the non-coding regions flanking the start and stop codons of genes annotated in genomic DNA, we see the expected sharp increase in dN/dS, consistent with well-defined coding sequence boundaries (Fig. S2B).

Second, the alignment of multiple homologous CDS is also error prone (*1*, *2*), and although this has not been documented, errors accumulating at the edges of the MSA could lead to increased measures of substitution rates owing to misaligned codons. We computed 16,590 simulated MSA of synthetic CDS, considered as ‘true’ alignments, and then realigned the synthetic CDS using several algorithms commonly used in the field, including MAFFT(*3*) used for Figure 1. We show that MSA algorithms do not create substitution biases along synthetic CDS length (Fig. S2C). Finally, because forces such as GC-biased gene conversion have been shown to affect molecular evolutionary rates (*4*, *5*), we examined the nucleotide and amino-acid composition of CDS and protein sequences for biases that could potentially explain the observations. Amino acid compositions are remarkably stable across protein length (Fig. S2D). Increase or decrease in frequency towards the N-terminus are visible but in magnitudes that are negligible compared to the 2-fold excess in dN observed here (Fig. 1D). While GC levels show a notable excess towards the 5’ end of the CDS (Fig. S2E), as previously shown, here too, neither the pattern nor the magnitude are compatible with the increase in dN/dS that we observe. In addition, GC-biased gene conversion is effectively neutral and would primarily affect the dS, which we show here to be constant throughout the length of the protein. Our observations are therefore robust to controls designed to address possible technical artefacts in the process from CDS annotation to substitution calculations.

#### 2. Modelling dN/dS variation in protein sequences:

We consider that the selective pressure measured as dN/dS at a given average position in protein sequences is under the influence of two forces: the proportion $\delta$ of residues at this position that are included in functional domain and the proportion $\alpha$ with high solvent accessibility (RSA > 0.7). Note that $\delta$ and $\alpha$ cover largely independent sets of residues, as shown in Figure S4: only 12% of residues overlapping domains have an RSA > 0.7, and only 17% of residues with an RSA > 0.7 overlap domains. The average dN/dS at a given position in proteins is then given as:

$$\frac{dN}{dS} = \frac{\delta}{\delta+\alpha}\cdot\frac{d\hat{N}}{dS}(domains) + \frac{\alpha}{\delta+\alpha}\cdot\frac{d\hat{N}}{dS}(highaccessibility)$$

With $\frac{d\hat{N}}{dS}(domains)$ the average dN/dS inside domains and $\frac{d\hat{N}}{dS}(highaccessibility)$ the average dN/dS in high solvent accessibility residues. The model was applied to the 7,614 human protein sequences present both in the AlphaFold set (for RSA computation) and in Ensembl. In this set, $\frac{d\hat{N}}{dS}(domains)$ = 0.14 and $\frac{d\hat{N}}{dS}(highaccessibility)$= 0.32. The mean absolute error (MAE) is computed as the mean of the differences between the model and the true value at each position. The percentage difference is computed as the ratio between the MAE and the dN/dS given by the model. When averaged across the three 50-codons panels the MAE is dN/dS=0.041 and the mean percent difference is 5.9%.
The model was also applied to a set of human protein without signal peptides, to see if this would improve the fit in the N-terminus region where it appears asymmetrical compared to the C-terminus in Figure 2E. We predicted signal peptides in our set of human protein sequences using SignalP version 5.0 (*6*) with default parameters, resulting in 15.82% proteins with a signal peptide (Fig. S5B).

#### Material and Methods

##### 1. Coding sequences, protein sequences and alignments

*(See Table S1 below for a summary of data set size (number of sequences) related to figures.)*

The sequence data comprises 4 sets covering different taxonomic groups: primates, rodents, plants, and Human-Mouse orthologs.

Primate and Rodent sequences were downloaded from the Ensembl database (*7*) as follows:

***Primates***. The genomes are from the following species: *Otolemur garnettii, Microcebus murinus, Propithecus coquereli, Prolemur simus, Saimiri boliviensis boliviensis, Cebus capucinus, Aotus nancymaae, Cercocebus atys, Mandrillus leucophaeus, Papio anubis, Theropithecus gelada, Macaca mulatta, Macaca fascicularis, Macaca nemestrina, Chlorocebus sabaeus, Colobus angolensis palliatus, Piliocolobus tephrosceles, Rhinopithecus roxellana, Rhinopithecus bieti, Pongo abelii, Gorilla gorilla, Pan troglodytes, Pan paniscus, Homo sapiens, Nomascus leucogenys, Carlito syrichta, Mus musculus* (outgroup).

**Rodents**. The genomes are from the following species: *Tupaia belangeri, , Dipodomys ordii, Jaculus jaculus, Rattus norvegicus, Mus musculus, Mus spicilegus, Microtus ochrogaster, Cricetulus griseus, Mesocricetus auratus, Peromyscus maniculatus bairdii, Nannospalax galili, Octodon degus, Cavia porcellus, Chinchilla lanigera, Sciurus vulgaris, Marmota marmota*

*marmota, Urocitellus parryii, Ictidomys tridecemlineatus, Ochotona princeps (outgroup), Oryctolagus cuniculus (outgroup).*

Phylogenetic gene trees and CDS sequences restricted to either primate or rodent genomes were downloaded from Ensembl Multi compara v101 (primates) and v104 (rodents) via the Perl API. To retain only strict 1:1 orthologs, for each tree the largest sub-tree that does not contain duplications was extracted, with a random choice in case of a tie. To avoid alignments with too few sequences, only the trees of size greater than 4 were selected. A multiple sequence alignment (MSA) on the amino acid sequences was then performed with MAFFT (*3*) (--maxiterate 1000 –localpair). Sequences were finally back-translated (treebest backtrans -t 0.9) with the corresponding CDS to obtain the final aligned codons in nucleotides.

**Plants.** The genomes were from the following *Fabids* species: *Cucumis sativus, Medicago truncatula, Lotus japonicus, Glycine max, Phaseolus vulgaris, Phaseolus angularis, Lupinus angustifolius, Manihot esculenta, Populus trichocarpa, Prunus persica.*

Coding sequences for all 10 genomes were downloaded from the April 2021 release of the OMA database (*8*) based on Ensembl Plants. A MSA on the amino acid sequences of each OMA family was then computed with FSA (*9*) with default parameters. Sequences were finally back-translated (treebest backtrans -t 0.9) with the corresponding CDS to obtain the final aligned codons in nucleotides.

**Human-Mouse orthologs.** Gene CDS were directly extracted from mRNA sequences (transcripts) downloaded from the October 2020 release of AniProtDB (*10*). A reciprocal blastp was performed (e-value $1.10^{-3}$) to select orthologs. Pairs of matching sequences were then aligned using the Needleman-Wunsh algorithm (needle, whith -gapopen 10.0 -gapextend 0.5) from the EMBOSS package.

**Gap-less alignments.** Algorithms introduce gaps in MSA to accommodate insertion or deletions of amino acids. They are more frequent at the edges than in the middle of CDS, and they may bias the counting of substitutions in these regions: for every gap, there is one less site available to count substitutions in a column of the alignment, thus decreasing statistical power and increasing noise. To quantify substitutions in an unbiased way, we removed, in each alignment, sequences that introduce gaps. Remaining sequences therefore are aligned with no gaps, only substitutions. Since it was not always possible to find at least 2 remaining sequences in a given alignment, this procedure resulted in a smaller set of alignments (see table 1 below).

| Group | Origin | Original Download | 1:1 orthologs | Gap-less | Present in Ensembl |
|---|---|---|---|---|---|
| Primates | Ensembl v101 | 20,275 | 16,810<br>*Figure 1A* | 16,248<br>*Figure 1B, 1D*<br>*Figure 2B* | NA |
| Rodents | Ensembl v104 | 17,774 | 13,668<br>Figure 3B, 3C | 12,170<br>*Figure 3B* | NA |
| Plants | Oma<br>(April 2021) | 33,425 | 33,095 | 7,513<br>*Figure 1C* | NA |
| Human-Mouse | AniProtDB | 35,045 Human<br>37,064 Mouse | 10,390 | 2,747<br>*Figure S2A* | NA |
| AlphaFold Human | AlphaFold Protein Structure Database | 23,391<br>*Figure 2C* | NA | NA | 7,614<br>*Figure 2D* |
| AlphaFold Mouse | AlphaFold Protein Structure Database | 21,615 | NA | NA | 5,893 |

**Table S1.** Number of protein sequences and gene families used in the analyses, and corresponding figures.

**Position-specific codon alignments and evolutionary rate computation**

To compute molecular rate presented in 3 panels of 50 codons each, the following procedure was followed. Two random CDS sequences beginning with a start codon ATG were chosen in each MSA, and the aligned codons at the same position in each pair were concatenated into a new, position-specific alignment. The procedure was applied for the first, middle and last 50 codons of each alignment. On each concatenated position-specific alignment, the dN and dS were computed using the YN00 model in Codeml from the PAML4 package (*11*) using the Bio.Phylo.PAML.codeml library. See Figure S1 for a schematic representation of the procedure. Several codon models were tested (YN00, LWL85 and NG86), with similar results.

**Metagenes**

Representations scaled from 0% to 100% of the length of proteins (metagenes) are computed as follows. For each MSA (e.g. for evolutionary rates, Figure 1A) or each protein sequence (e.g. domain density, Figure 3A), the length of the MSA (resp. sequences) is divided in 100 intervals, the variable of interest for each interval is added to an interval-specific total across all MSA (resp. sequences), and the average value for that interval is plotted on the Y-axis.

**Synthetic sequences**

To control for the potential role of errors during MSA constructions, which may have caused the observed excess of substitutions in CDS edges, we built a set of 16,590 synthetic multiple alignments using INDELible (*12*), which simulates aligned sequences containing insertions/deletions. Parameters are LAV 2 300 model for gaps, submodel 4 0.175 , insertrate 0.010500, deleterate 0.021000, and primates values for codons stationary frequencies. The template tree is taken from the real dataset, with one replicate for each gene tree. This results in 16,591 simulated alignments which are considered to be the "truth". All gaps are then removed to reconstitute the original CDS, which are translated before being aligned with mafft (--maxiterate 1000 --localpair) in amino acids and back-translated into codon alignments. There are thus two sets of 16,591 sequences: a "truth" set generated by INDELible and a "realigned" set. It is then possible to compute the dN/dS ratio for both datasets and compare the values at each position to measure the impact of the multiple alignment method. The same experiments have been performed by realigning with FSA with and without HMM-Cleaner (*13*), with similar results. Other experiments were conducted by artificially lengthening (by a factor of up to 50) the length of the branches to obtain more distant sequences, but this did not change the conclusions.

**Flanking regions**

To control for a potential bias due to annotation errors, particularly at the extremities of the coding sequences which are notoriously difficult to identify precisely, we computed dN/dS values on position-specific codon alignments that included Untranslated (UTR) sequences before the start codon and after the stop codon. For this, all *Nomascus leucogenys* (Nleu 3.0) and *Pan troglodytes* (Pan tro 3.0) CDS sequences from Ensembl version 104 were downloaded with upstream and downstream flanking 30 nucleotides collected via BioMart. Amino acid translation, double blastp (-evalue 1e-3) and reciprocal best hits selection followed by alignment (mafft --maxiterate 1000 --localpair) and finally backtranslation were performed to obtain aligned nucleotide data. Only the 13,344 pairs of orthologs aligned with no gaps were retained in order to compute dN/dS in these CDS as well as in the 10 flanking pseudo-codon overlapping UTR sequences.

**Functional Domain distribution**

Functional protein domains were downloaded via the Ensembl v101 API for 12,067 human protein sequences selected from the primate MSA for the computation of molecular rates, for four different databases: Pfam (*14*), Smart (*15*), SuperFamily (*16*) and Prosite (*17*) patterns. To compute random re-distribution of domains, the sizes of the inter-domain spaces for each protein were summed, then redivided in the same number of intervals but using randomly sampled boundaries. These new inter-domain intervals were then re-inserted between the original domains without changing their order or content.

**Solvent accessibility**

PDB-formatted 3D structures of the human and mouse proteomes predicted by AlphaFold (*18*) were downloaded from the AlphaFold Protein Structure Database (*19*) resulting in 23,391 human and 21,615 mouse structures. To obtain relative solvent accessibilities (RSA), we used pCASA (*20*) to compute the total accessible surface (m) and the average accessible surface (n) for each residue, and took the ratio between the former and the latter: RSA = m / (n + 18). The average accessibility was incremented by 18, a constant accounting for the part of each residue that is included in the peptidic chain and almost never accessible. Finally, the exact intersection of the protein set available from the AlphaFold database and of the protein set used in our study to compute molecular rates resulted in 7,614 human and 5,893 mouse proteins.

**Signal Peptides**

Signal peptides were identified in the 16,248 primate sequences involved in gap-less multiple sequence alignments (Table S1) using SignalP (*6*) resulting in 15.82% proteins with a signal peptide.

**Pearson correlations in individual sequences**

We started from the same 13,668 rodent phylogenetic trees and sequence families as described above from Ensembl version 104 (Table S1), aligned with MAFFT as described. To maintain a balance between statistical power and robustness dN/dS estimates, we restricted the alignments to those containing sequences from at least five different species, where the first 50 and last 50 codons columns have at least 10 columns that have a substitution yet where gaps are a minority of columns. A Pearson coefficient from a linear correlation was computed for each alignment between the dN/dS values obtained at each 50 codons and their respective positions in these 50 codons

**Positive selection**

To estimate sites under positive selection, we used 13,668 rodent phylogenetic trees and sequence families as described above from Ensembl version 104 (Table S1), aligned with MAFFT as described (Fig. 3B and 3C). To compute positive selection in high and low RSA amino-acids, we had to restrict the analysis to alignments for which we could unambiguously find a correspondence between the AlphaFold dataset and the Ensembl Dataset, thus ending with 5,893 MSA. We used Hyphy MEME (Mixed Effects Model of Evolution) (*21*, *22*) to detect sites evolving under positive selection under a proportion of branches. In all cases, only positions with a positive selection p-value lower than 0.05 were retained.

We also collected data from the literature (Figure S6) as follows:

- Table S5 in supplementary data in van der Lee et al. (2017) (*23*), corresponding to 934 Positively Selected Residues (PSR) in 331 primate protein coding genes out of 11,096 initial gene families tested. Positively selected genes were identified by Codeml from the PAML package (*11*) using a gene-level dN/dS test and positively selected residues by selecting sites with a significant Bayesian posterior probability.
- Tables S4-S5-S7-S15-S16-S18 in Murrell et al. (2012) (*22*): We retained only 13 positively selected sites identified jointly using HyPhy-MEME and FEL (M+F+) in 7 animal genes.
- Table 1 in Rodrigue et al., (2021) (*24*), corresponding to 51 positively selected sites identified in 6 metazoan genes with the MutSel-M3 model at a threshold $p > 0.95$.
- Table 1 from Yokoyama, S. (2008) (*25*) Corresponding to 51 experimentally validated sites in 6 visual pigment proteins (e.g. Rhodopsins).

#### Supplementary figures

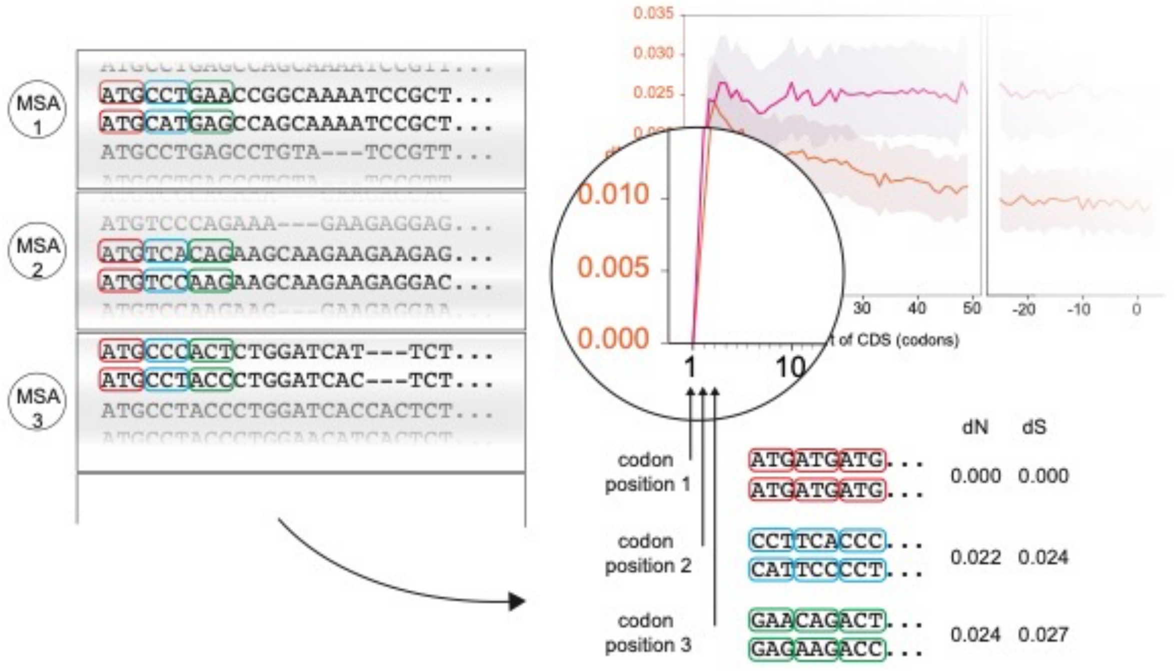


**Figure S1.** Schematic representation of the strategy used to compute position-specific average dN and dS in coding sequences. Starting from several thousand multiple sequence alignments (MSA), 2 sequences from each are randomly chosen. In each pair, aligned codons are extracted and concatenated into new, position-specific alignments. In the example given, the first codons (always ATG coding for methionine, in red) are concatenated into a new Codon1-specific alignment consisting only of ATG codons in both upper and lower sequence, thus resulting in dN = dS= 0.0. The second codon is variable (in blue) among MSAs, and once the thousands of 2$^{nd}$ codons are concatenated, they provide an average dN and dS value for codons at the 2$^{nd}$ position of coding sequences.

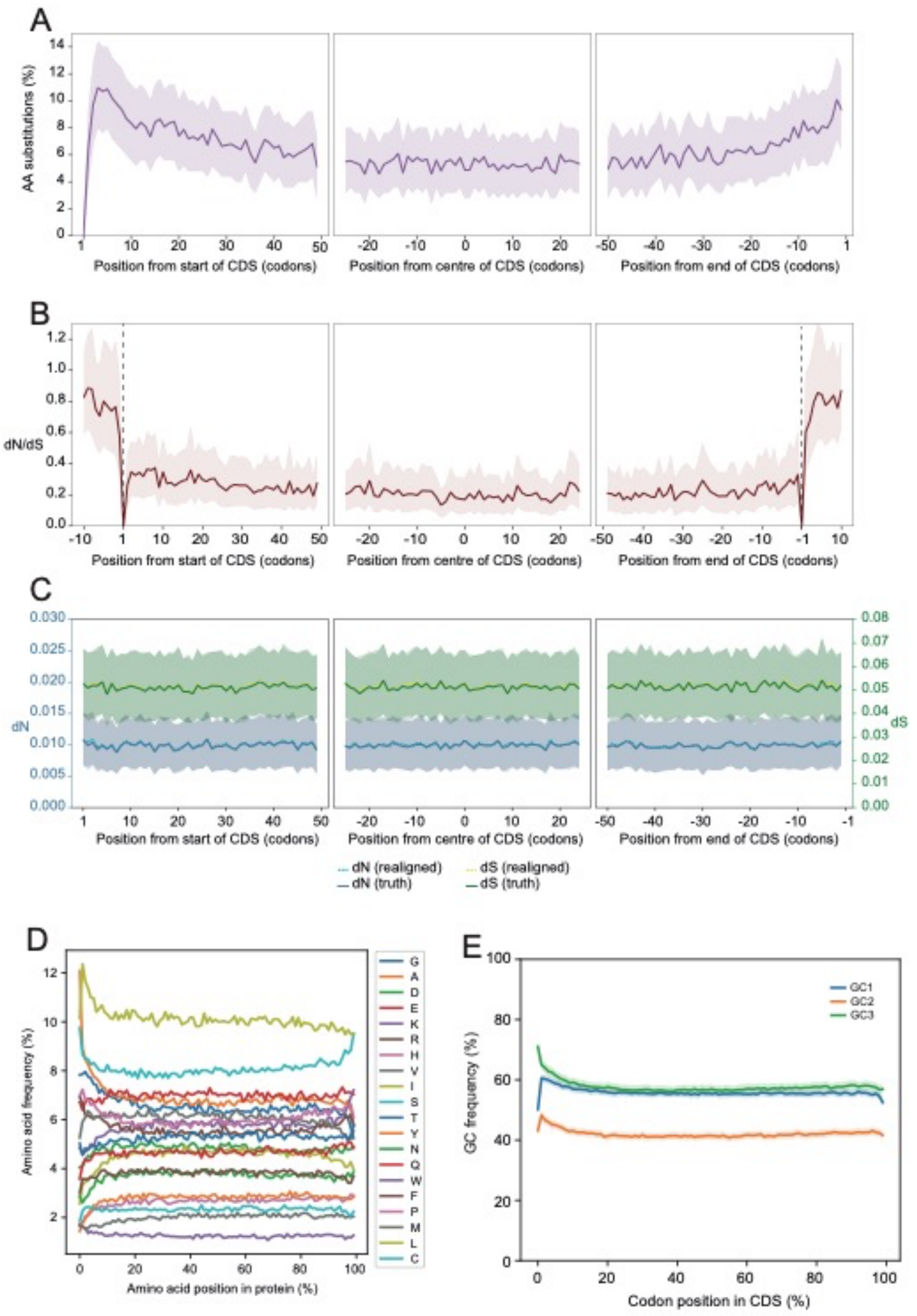


**Fig. S2**: (**A**). Distribution of amino acid substitution in 2,747 alignment of protein sequences directly inferred from aligned human and mouse transcript sequences. The shaded area is the 95 CI computed from 1000 samplings of 1000 alignments (**B**) dN/dS computed from the CDS of 13,344 pairs of primate sequences (*N. Leucogenys* and *P. Troglodytes*) including 30 bases (10 triplets) flanking the start and stop codons. (**C**) dN and dS computed on 16,591 CDS simulated and aligned with INDELible (truth, plain lines), and after being realigned using MAFFT (realigned, dashed lines). (**D**) Amino acid frequency distribution along 296,433 predicted protein sequences from 26 primates proteomes downloaded from the Ensembl database version 101, shown on a scale of 0-100% of the protein length. (**E**) Guanine and Cytosine (GC) frequency distribution along 296,433 CDS from 26 primates as in A. The GC% distinguishes the first (GC1) the second (GC2) and the third (GC3) codon position. Shaded areas are the distribution of GC% obtained from 1000 resamplings of 1000 gene families.

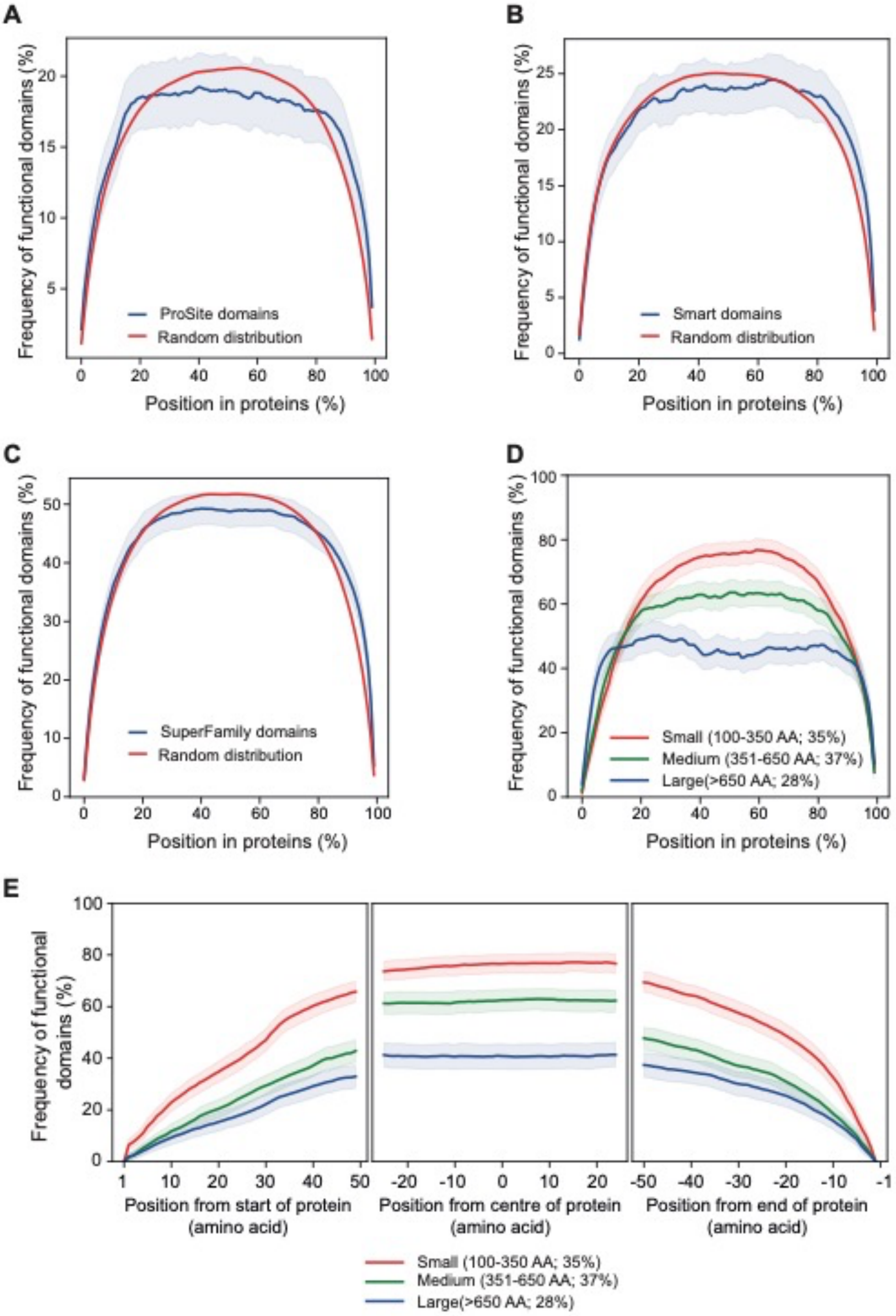


**Figure S3**. Distribution of functional domains in proteins as predicted by Prosite (**A**), Smart (**B**) and SuperFamily (**C**). Distributions are shown for 12,067 human proteins from Ensembl v101, along their length rescaled from 0-100%. In each case, the true distribution is shown in blue, and the random distribution is shown in red for comparison. The blue shaded area is obtained by 1000 resamplings of 1000 protein sequences. (**D**) Distribution of PFAM functional domains in 12,067 human protein sequences classified in small, medium and large protein length categories, showing that protein termini in small proteins exert a stronger influence on domain distribution. (**E**) Same as in D but in absolute amino-acid coordinates in protein sequences, shown here for the first, middle and last 50 amino-acids.

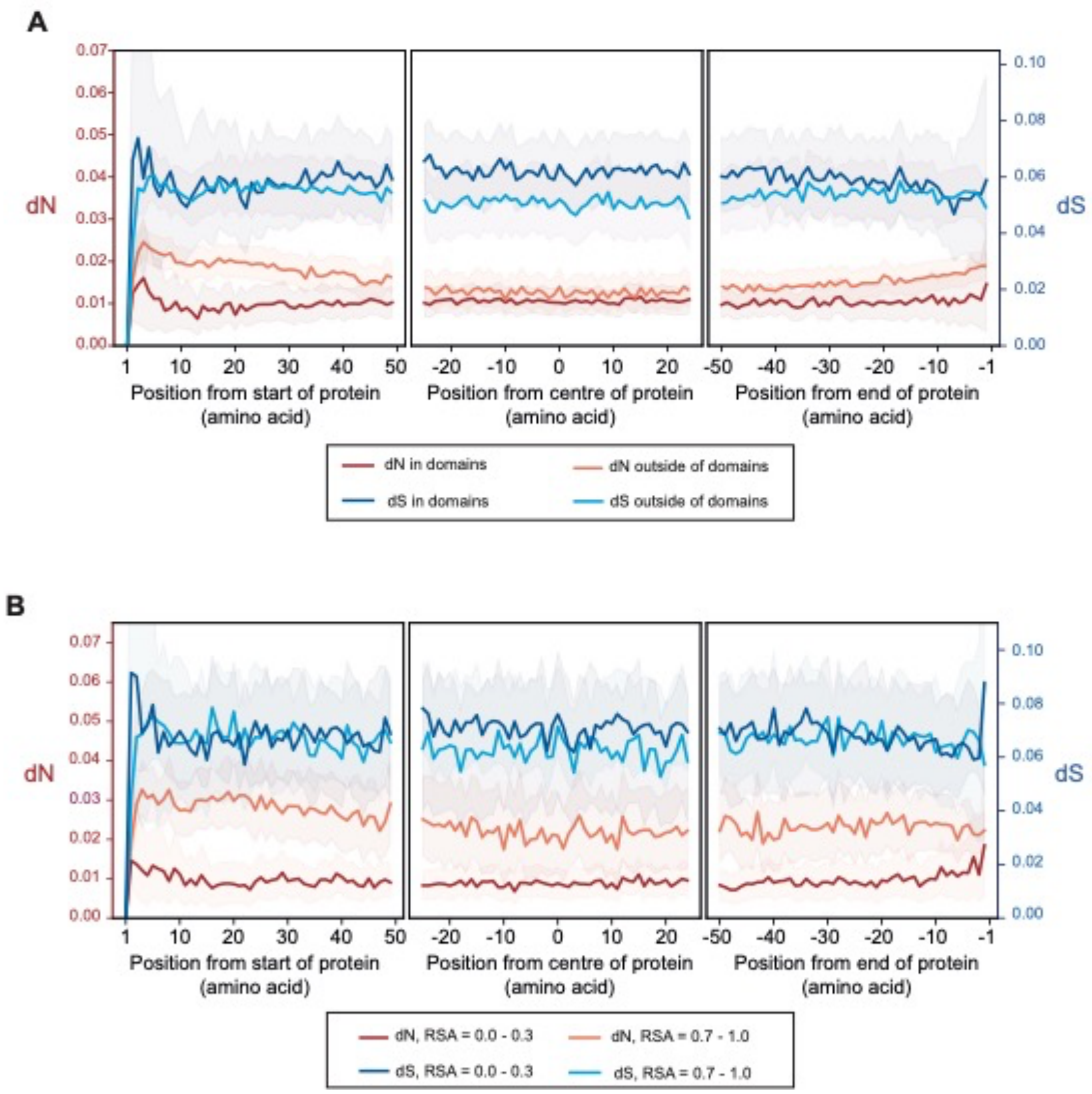


**Figure S4**. (**A**) Distribution of dN and dS inside and outside of domains. (**B**) Distribution of dN and dS in low (RSA < 0.3) and high (RSA > 0.7) relative solvent accessibility amino acids.

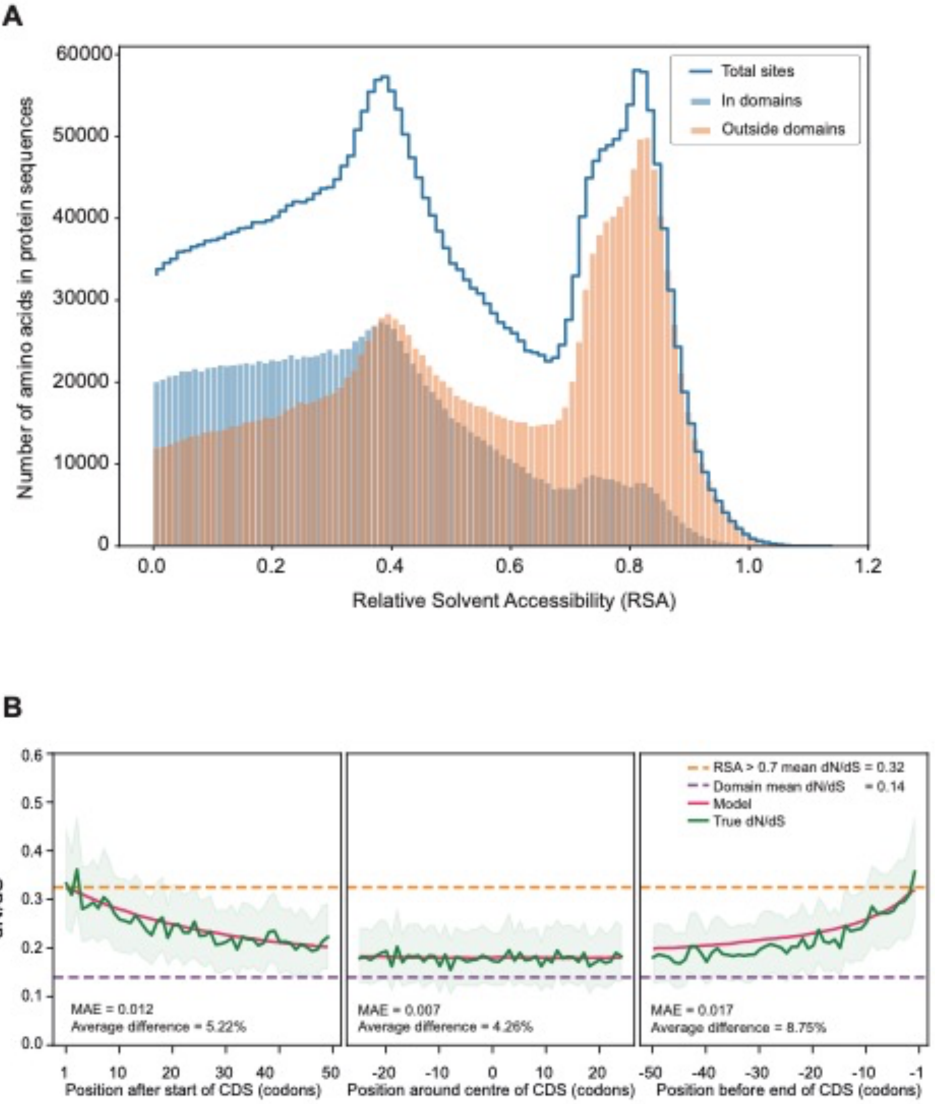


**Figure S5**. **(A)** Distribution of amino acid counts in 8,843 human protein sequences present in the AlphaFold dataset and in our dataset, inside and outside PFAM domains, as a function of their Relative Solvent Accessibility (RSA).(**B**) Model of dN/dS variation (red) as a function of the proportion of sites with high RSA and in functional domains (as in Figure 3E), but excluding proteins with a predicted signal peptide.

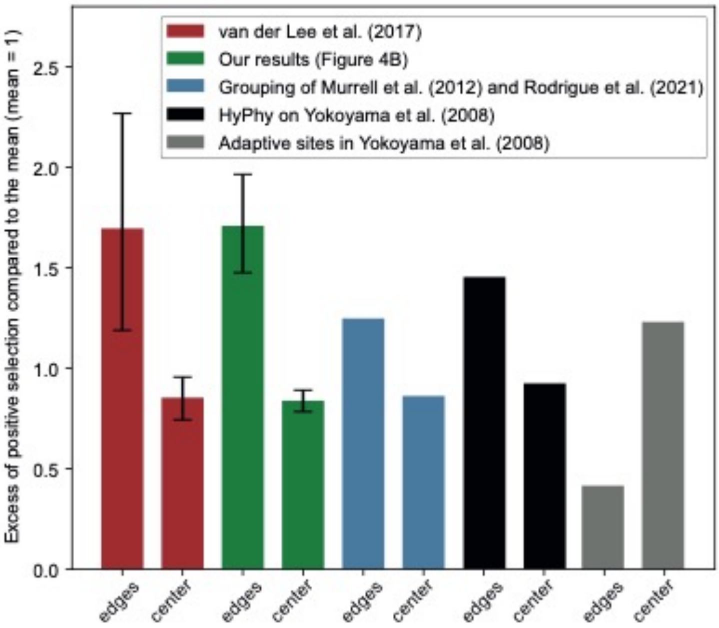


**Figure S6**. Estimation of positive selection by computational and experimental methods from the literature and this study. See Methods for details on each dataset. "Edges" represent the sum of sites identified under positively selection in the first and last 50 codons of the proteins, while "center" represent the sites detected in the central 50 codons. For the large datasets (van der Lee, n= 934 sites; our results n= 7,261 sites) we performed a bootstrap by resampling 1000 times 50 sites (van der Lee (2017)) or 1000 sites (our data), and the interval shows the 95% confidence interval.
